# Predicting Clinical Phenotypes by Growth Curve Modeling of Transcriptomic Signatures during Disease Progression

**DOI:** 10.64898/2026.01.13.699292

**Authors:** Mahboube Akhlaghi, Erfan Ghasemi, Meghana S. Ray, Saumyadipta Pyne

**Author notes:** Corresponding author: S. Pyne.

## Abstract

High-throughput transcriptomic analysis has benefited from many statistical tests of differential gene expression across two or more groups such as *t* tests, ANOVA, etc. Yet, in complex transcriptomic datasets such as multi-group longitudinal measures, few studies have addressed such key issues as group effects and temporal dependency in expression profiles with a single model that is both practically effective and theoretically grounded. In this study, we used Growth Curve Model (GCM), as a generalization of MANOVA, to identify differentially expressed longitudinal profiles of genes, and thus predicted the associated clinical phenotypes, of pediatric lupus during the progressions of the disease across two different racial groups. In particular, we detected a module of histone genes which was shown to be linked with lupus.

## 1. Background

The capability for prediction of an individual’s clinical phenotypes during their disease progression can offer a significant advantage to the clinician to not only be prepared for what might occur but also potentially treat those phenotypes in an effective and timely manner. In fact, new AI-driven applications have been developed to analyze disease dynamics and enable disease surveillance to predict specific phenotypes, and their corresponding medical codes, for future clinical visits of a given individual.(Choi *et al*., 2016; Shankar *et al*., 2023) Other platforms such as the Danish Disease Trajectory Browser use statistical analysis of population-scale medical data to forecast disease trajectories.(Siggaard *et al*., 2020)

Understanding a disease’s trajectory could be further complicated by its phenotypic heterogeneity, which makes it difficult to define the disease precisely in terms of a specific set of clinically-diagnosable phenotypes. This problem is particularly acute for certain autoimmune diseases, e.g., systemic lupus erythematosus (SLE). SLE is manifested as a chronic condition that can present several clinical phenotypes in a given individual, which can affect **–** either singly or together **–** multiple parts of the body, and possibly at different points of time occurring over many years, including as occasional severe “flares” of the disease. SLE is particularly heterogeneous in terms of the variety as well as the dynamics of its potential phenotypes, the diagnosis of which depends on both clinical manifestations (e.g., malar rash or organ damage) and laboratory tests (e.g., autoantibody detection). These phenotypes, in turn, vary by the underlying mechanisms-of-action and individual genetics. As a result, there is no current consensus among the various clinical criteria (e.g., ACR, BILAG, SLICC, SLEDAI) that are used for SLE diagnosis. For a detailed survey of the topic, see Dai *et al*. (2025).

To address the challenge of clinical phenotypic heterogeneity in disease dynamics, researchers are increasingly focusing on the molecular signatures underlying these phenotypes. In this direction, high-throughput gene expression data was found to be particularly useful for the prediction of phenotypes as outcomes of transcriptomic signals (Ellis *et al*., 2018). However, such data are generally not available in the form of longitudinal observations, which makes it difficult to model how differential gene expressions underlying the outcomes change during disease progression. Further, such phenotypes are known to evolve differently across different races and ethnic groups, and thus, the transcriptomic signals must be identified with a robust statistical model that incorporates such temporal dependencies and group effects.

SLE, commonly referred to as “lupus”, is most often diagnosed in young adults and is also disproportionately prevalent among women. In the United States (U.S.), it ranks among the top 10 causes of death for Black/African-American and Hispanic women of childbearing age, particularly between 15 and 45 years. In fact, 41% of all estimated SLE cases in the U.S. are Black (along with Native American and Asian) and 21% are Hispanic (Feldman *et al*., 2013). According to the Lupus Foundation of America (lupus.org/health-disparities), the prevalence of SLE is thus 3 and 2 times higher in Black and Hispanic women respectively compared to their white counterparts. Such racial disparity has led to recent studies of ethnicity-specific molecular signatures of SLE, e.g., Andreoletti *et al*. (2021).

In this study, our objective is to perform a systematic analysis of longitudinal high-throughput gene expression data to identify differential gene expression signatures in female cohorts of pediatric lupus in black and Hispanic cases during the course of their disease progression. The study is balanced with an equal number of temporal observations made per group, and both groups comprise of participants in the same age range. Further, the statistically significant longitudinal transcriptomic signatures were tested against well-known associations with curated clinical phenotypes to identify collections of distinct outcomes of pediatric lupus across the two most seriously SLE-affected population-groups in the U.S.

Towards this aim, we developed a novel framework for integration of 3 components. First, we used longitudinal high-dimensional gene expression data from a clinical study (Banchereau *et al*., 2016a) to detect longitudinal transcriptomic signatures that represent the molecular dynamics of SLE over time across 2 racial-groups of female pediatric cases. Second, the framework leverages on an established and curated database named ‘Human Phenotype Ontology’ (HPO) for its annotations of hundreds of known clinical phenotypes in terms of the human genes that are associated with each of them.(Gargano *et al*., 2024) Third, we used Growth Curve Models (GCM) as a powerful and general multivariate approach that was shown to be effective in identification of differential patterns of longitudinal gene expression across different groups of individuals (Jana *et al*., 2017; Hamid and Beyene, 2009).

The clinical data used in this study are in the form of short time series of longitudinal gene expression measurements for each individual who belongs to one of the *k* = 2 balanced groups that were compared. While the measurements across individuals are assumed to be independent, those within an individual may be temporally correlated. In the following section, we provide a brief review of the well-known GCM approach leading up to its application in high-dimensional “omic” data analysis. The Methods section describes both the analytical pipeline as well as the details of the SLE longitudinal gene expression dataset used in this study. The Results section identifies the significant longitudinally differentially expressed genes including a detected module of histones, and the corresponding clinical phenotypes. Finally, we end with a Discussion of the model and its application in this study.

## 2. Background of GCM

The analysis of growth curves began with a simple yet critical statistical challenge: how do we efficiently compare relatively short yet complex profiles of correlated observations made over time? The solution demanded methods that could reduce the data’s dimensionality without sacrificing the biological truth (Rao, 1958).

### 2.1. An Efficient Model for Summarizing Growth

Early work, notably by Wishart (1938), recognized that collecting observations on a growing organism, either continuously or at specified time points, results in correlated data (Box, 1950; Leech and Healy, 1959; Rao, 1959; Elston and Grizzle, 1962; Bock, 1963). Wishart fitted a second-degree polynomial to individual growth curves, summarizing many observations with just a few coefficients (linear and quadratic terms) (Wishart, 1938).

While the classic paper by Potthoff and Roy (1964) is generally considered to be the first paper in growth curve modeling (see below), a similar model was introduced earlier by C.R. Rao (1958). Following Wishart, Rao emphasized on “replacing the various observations on growth by a few summary figures which lead to most efficient comparisons between groups”, particularly when facing small samples and large variations between individual curves. He explored simplifying complex growth functions using a time metameter *τ* = *G*(*t*), which could transform the function into a simple linear rate. If time is transformed using this function *G*(*t*) such that the growth rate becomes uniform with respect to the chosen time metric, then the process can be adequately represented in terms of the initial value and a redefined constant growth rate. This allowed the entire set of observations to be summarized by the initial value (**Y**_0_) and a single estimated rate(**b**). This estimated rate **b** was calculated similarly to a regression estimate, proportional to the weighted sum of gains (**y**_*i*_)(Rao, 1958):

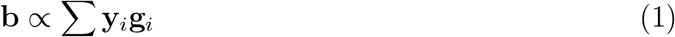

Here, the length of the interval between the (*i −* 1)-th and *i*-th time points on the transformed time axis is represented by *g*_*i*_.

This approach provided a valid test for the null hypothesis that the average growth curve is the same across different treatment conditions, irrespective of assumptions regarding the curve’s specific nature(Rao, 1958).

### 2.2. Standardizing the GCM Framework

The mathematical structure for advanced growth curve analysis evolved from the generalized application of linear hypotheses within multivariate analysis. Rao (1959) formalized the analysis for *p* correlated normal variables (**y**) where the expected mean vector *E*(**y**) is linearly related to *m* unknown parameters *τ* :

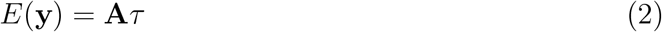

This framework allowed for rigorous testing of model adequacy, such as whether an average growth curve could be represented by a polynomial of a given degree (Hsu, 1938; Rao, 1959). **A** is the matrix of coefficients which is known. The formalization led directly to the Generalized MANOVA (GMANOVA) model, also known as the Growth Curve Model (GCM) (Khatri, 1966; Kollo and von Rosen, 2005).

#### A. Formal Model Definition

The GCM introduced by Potthoff and Roy (1964) models the expected mean using a bilinear structure that incorporates both group effects and the time structure (Hamid and Beyene, 2009; Jana *et al*., 2017). Consider a study with *k* groups, where each individual has repeated measurements taken at *p* time points. In the *i*-th group, there are *n*_*i*_ individuals (*i* = 1, 2, … , *k*), resulting in a total of 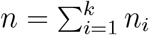 individuals across all groups.

If we consider that the expected response for the *i*-th group changes over time according to a polynomial function of time *t* with degree *q −* 1:

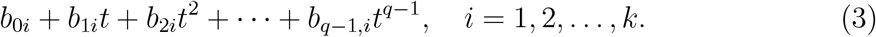

This setup defines the Growth Curve Model (GCM), which can be expressed in matrix form as:

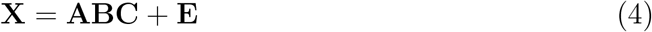

Here, **X** (*p × n*) is the observation or response matrix, **A** (*p × q*) is the within-individual design matrix incorporating temporal ordering and time points as a continuous variable, **B** (*q × k*) is the unknown parameter matrix containing the coefficients of the polynomial growth curves, **C** (*k × n*) is the between-individual design matrix for group effects, and **E** (*p × n*) is the error matrix (Jana *et al*., 2020).

While **A** and **C** are known matrices, **B** is an unknown parameter matrix consisting of the coefficients of the polynomials described in Equation 4. The between-individual design matrix **C** is the same design matrix as in univariate and classical multivariate linear models. The columns of **E** are assumed to be *p*-variate independently normally distributed random vectors with mean zero and an unknown positive definite covariance matrix **Σ**.

In the early formulation of the Growth Curve Model (GCM), one common objective was to compare the mean trajectories of two biological groups. Specifically, consider two groups where the aim is to assess whether the average expression pattern of a given gene *g* is the same across both. The hypothesis can be expressed as:

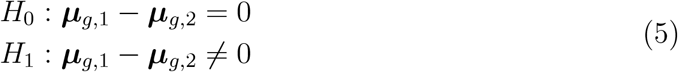

Here, ***μ***_*g*,1_ and ***μ***_*g*,2_ represent the average expression levels of gene *g* under the two biological conditions, respectively. An equivalent formulation can be written in terms of the parameter matrices:

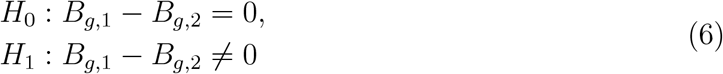

Such hypotheses within the GCM framework were first introduced by Potthoff and Roy (1964), who provided the foundational statistical structure for comparing group mean profiles over time (Hamid and Beyene, 2009).

Potthoff and Roy (1964) showed that the GCM problem could be reduced to a standard MANOVA problem via a transformation matrix involving an arbitrary symmetric positive definite matrix **G**. They provided an estimator for **B**:

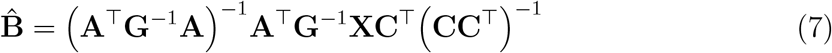

This reliance on an arbitrary choice of **G** was noted to affect the variance of the estimators and drew criticism.

#### B. Classical Maximum Likelihood Estimation

Khatri (1966) derived the definitive Maximum Likelihood Estimators (MLEs) for **B** and the covariance matrix **Σ**, assuming multivariate normal errors and full rank design matrices. The MLE for **B** solved the problem of arbitrary **G** by replacing it with the estimated Sum of Squares and Products (SSP) error matrix **S**:

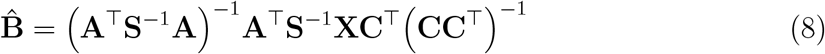

where

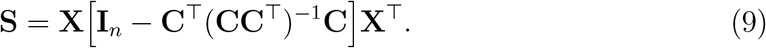

This 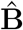 estimator is unique under full rank conditions and agrees with Potthoff and

Roy’s estimator when **G** = **S**.

For further details, the interested reader is referred to the excellent texts on the topic that were published both in the past (Srivastava and Khatri, 1979; Kshirsagar and Smith, 1995; Pan and Fang, 2002; Kollo and von Rosen, 2005) as well as more recently (von Rosen, 2018).

### 2.3. Addressing High Dimensionality of Genomic Data

Applying GCM to modern time course microarray data involves a specific statistical obstacle: gene-specific high-dimensionality (*n < p*). Since the sample size (*n*) is smaller than the number of time points (*p*), the sample covariance matrix (**S**) becomes singular, invalidating traditional MLE methods requiring **S**^*−*1^ (Tai and Speed, 2006).

Hamid and Beyene (2009) addressed this by proposing a GCM approach that explicitly uses time dependence and handles singularity via moderation. The gene-specific GCM remains **X**_*g*_ = **AB**_*g*_**C** + **E**_*g*_ (where **A** and **C** are design matrices common to all genes). To solve the singularity issue, generalized inverse techniques were adopted. The conventional MLE for **B** is modified by replacing the inverse of the SSP matrix **S** with the Moore-Penrose generalized inverse(**S**^*−*^) (Moore, 1920; Penrose, 1955). This yields the moderated Maximum Likelihood Estimator 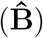:

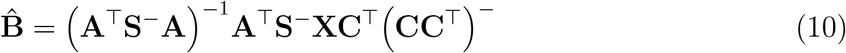

This generalized inverse technique was also used to derive a moderated trace test statistic, ensuring the test remained applicable even in high-dimensional scenarios where *n < p*.

Alternatively, the singularity problem can be tackled using moderation techniques like the Empirical Bayes approach (Lönnstedt and Speed, 2002; Smyth, 2004; Cui *et al*., 2005). This method utilizes a transformation **Λ**^*−*1^**A**^⊤^ **AΛ**^*−*1^**A**^⊤ *−*1^ (similar to Potthoff and Roy’s method) where the matrix **Λ** incorporates information borrowed across thousands of genes. This process results in a non-singular moderated covariance matrix 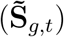 used in the moderated likelihood ratio test 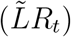 to rank genes (Tai and Speed, 2006):

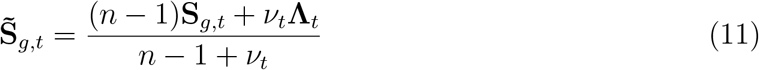

In this formula *v*_*t*_ *>* 0 and *v*_*t*_Λ_*t*_ *>* 0 are the degrees of freedom and the scale matrix for each treatment (*t*) condition, respectively and *g* represents different genes. The use of GCM (which incorporates temporal mean structure) for ranking genes was demonstrated to perform uniformly better than approaches ignoring this structure (classical MANOVA) when time dependence exists in the data (Hamid and Beyene, 2009).

## 3. Bioinformatic Analysis

### 3.1. Gene Expression Data

We utilized gene expression data from the GEO database (GSE65391), which was already preprocessed and normalized prior to download (Banchereau *et al*., 2016b). For consistency, we included only systemic lupus erythematosus (SLE) patients with available data for their first four clinical visits, and restricted our analysis to those visits. To ensure a balanced study design, we focused on two racial groups, African-American (AA) and Hispanic (H), each including exactly 11 and only female participants, all of them with SLE, and belonging to the same age-range of 11-20 years. For each participant, we have expression data of the same genes in the whole blood samples collected by the same study and measured using the same microarray platform across four clinical visits. Thus, we systematically compared gene expressions across female pediatric cases of two racial groups, AA and H, during the course of SLE progression.

The 22 samples used in our study are specified in GEO as “whole blood-SLE-#” such that the 11 AA Id #s are 95, 133, 152, 177, 212, 230, 231, 234, 260, 313, and 326; and the 11 H Id #s are 99, 125, 129, 178, 181, 211, 237, 264, 271, 304, and 337.

### 3.2. Methods

For preprocessing microarray data, probe-to-gene annotation was carried out using the `AnnotationDbi` R package (Pagès *et al*., 2025). Probes were mapped to human gene symbols, and in cases where multiple probes corresponded to the same gene, the median expression value was taken. Thus, we obtained longitudinal expression data for 14,797 genes.

To evaluate differential gene expression patterns over time between groups, we applied a Growth Curve Model (GCM) in a multivariate framework. This approach allowed us to test for group-by-time interaction effects. The key test statistic was a moderated trace statistic derived from the residual covariance structure:

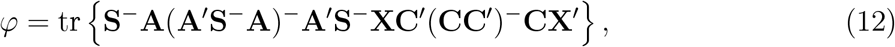

Where *X* denotes the matrix of gene expression observations, *A* is the design matrix for time points, *C* is the contrast matrix for group comparisons, *S* is the residual covariance matrix, tr represents the matrix trace, and ^*−*^ indicates the inverse or Moore-Penrose generalized inverse.

The residual covariance matrix *S* was calculated as:

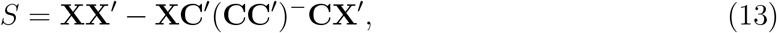

which represents residual variability after accounting for group effects. Depending on the relationship between the number of samples (*n*) and the number of parameters (*p*), we used either the standard inverse (*n > p*) or the Moore–Penrose generalized inverse (*n ≤ p*).

The design and contrast matrices used were:

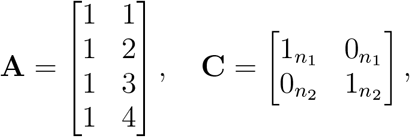

with the following parameters:

In this study, *K* = 2 represents the number of groups, *p* = 4 denotes the number of time points, *n* = 22 is the total number of patients, and *n*_1_ = 11, *n*_2_ = 11 indicate the number of patients per group.

To assess statistical significance, we performed a permutation test with 10,000 iterations (Onghena, 2017). Group labels were randomly shuffled across patients while preserving the longitudinal structure of repeated measures. For each permutation, the trace statistic was recalculated. A p-value was then computed as:

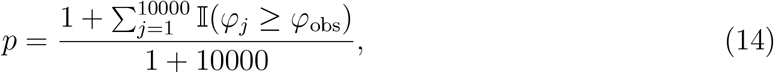

where *φ*_obs_ is the observed test statistic, *φ*_*j*_ is the statistic from the *j*-th permutation, and 𝕀 (·) is the indicator function. A p-value threshold of 0.05 was used to identify the significantly differentially expressed genes, which were shortlisted for downstream analysis.

Geneset over-representation analysis (ORA) is a widely used approach to test whether a finite set of genes (or geneset) is statistically over-represented in another geneset (Rivals *et al*., 2007). Typically, the first input geneset is an unordered list of genes of interest that is produced from bioinformatic analysis of a given dataset; and the second, also unordered, consists of genes involved in a known biological process or pathway that is usually selected from an annotated catalog of relevant genesets. A significant over-representation would suggest potential involvement of the former geneset (i.e., the list of significant genes from the above-mentioned trace test) in the latter (representing the various known SLE phenotypes and clinical outcomes, as described below).

The Human Phenotype Ontology (HPO) is a well-known database of standardized vocabulary of phenotypic abnormalities (each annotated with a unique HPO Id) found in human diseases (Robinson *et al*., 2008). We obtained a catalog of HPO genesets that are associated with different clinically observed SLE outcomes. Then ORA was performed using the `enricher` function from the `clusterProfiler` R package (Yu *et al*., 2012). It applied the hypergeometric test to evaluate whether our list of identified genes is significantly enriched in a HPO geneset relative to a background gene universe. The p-value from this over-representation test was calculated based on the hypergeometric distribution

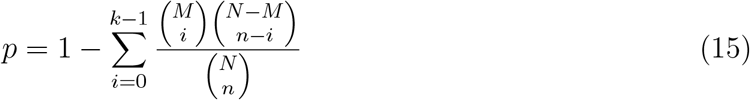

Here, *N* is the total number of genes in the background distribution, *n* the size of the first geneset, *M* the size of the second geneset, and *k* the number of genes in the first geneset that are annotated as members of the second geneset. The background distribution is, by default, of all annotated human genes. A p-value threshold of 0.05 is used for selecting the significant HPO genesets.

### 3.3. Results

Our dataset consists of the expression values of 14,797 genes measured in blood samples collected at 4 clinical visits during the progression of SLE in 22 pediatric cases from 2 racial groups (AA and H) of 11 each. Using the GCM-based trace test, we identified 720 genes that were significantly differentially expressed during the progression of SLE, and this list was used for ORA. Thus, we identified not only the gene expression profiles that distinguish the progression of SLE between the AA and H groups, but also the most significant SLE phenotypes, as per HPO, that are regulated by the same molecular signatures. Figure 1 shows these phenotypes beginning with mild proteinuria, a hallmark of lupus nephritis in which autoimmune attacks damage the kidney filters causing leakage of proteins into urine.

**Figure 1:**
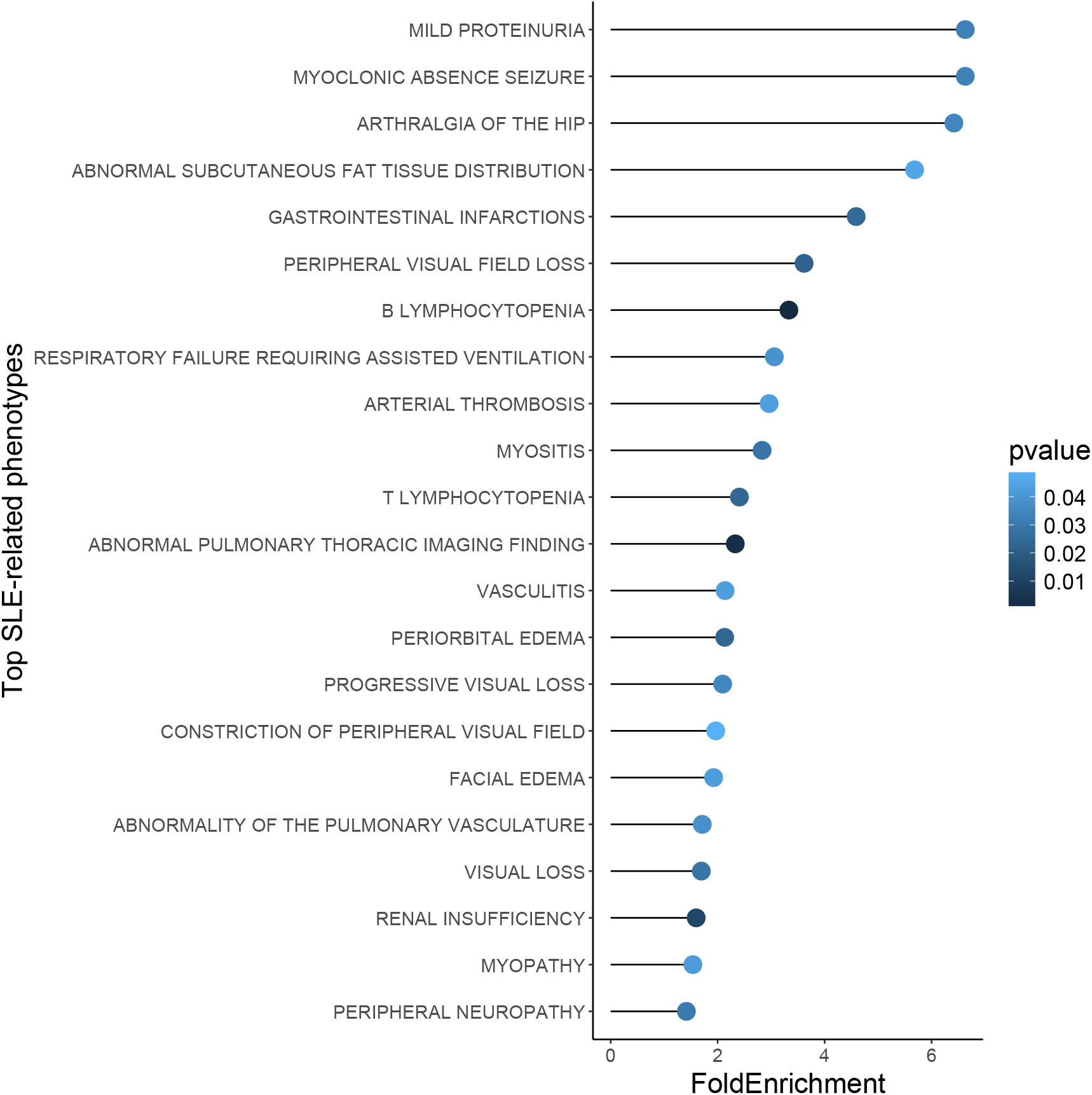
Over-representation of differentially expressed genes in the known SLE-related phenotypes as specified by the corresponding HPO genesets. The plot shows the top genesets from Over-representation analysis. Points represent fold enrichment for each pathway, with color indicating the corresponding p-value.

In particular, we identified a module of 9 histone genes that are known to be associated with SLE according to the curated Kyoto Encyclopedia of Genes and Genomes (KEGG) (Kanehisa and Goto, 2000). All of these histone genes showed marked differences in their mean longitudinal profiles across the 2 groups, as depicted in Figure 2. As indicated by their statistically highly significant p-values in Table 1, their mean longitudinal profiles remain consistently higher in AA than in H, and those differential patterns appear throughout the module as the disease progresses in Figure 2. The histones H2A, H2B, H3, and H4 are known to be involved in the KEGG pathway of neutrophil extracellular traps (NET) formation (Kanehisa and Goto, 2000). Increased NET accumulation and impaired clearance are associated with more severe disease, organ damage, and potential flares, making it a driver and a biomarker of pediatric lupus (Giaglis *et al*., 2016; Wigerblad and Kaplan, 2022).

**Figure 2:**
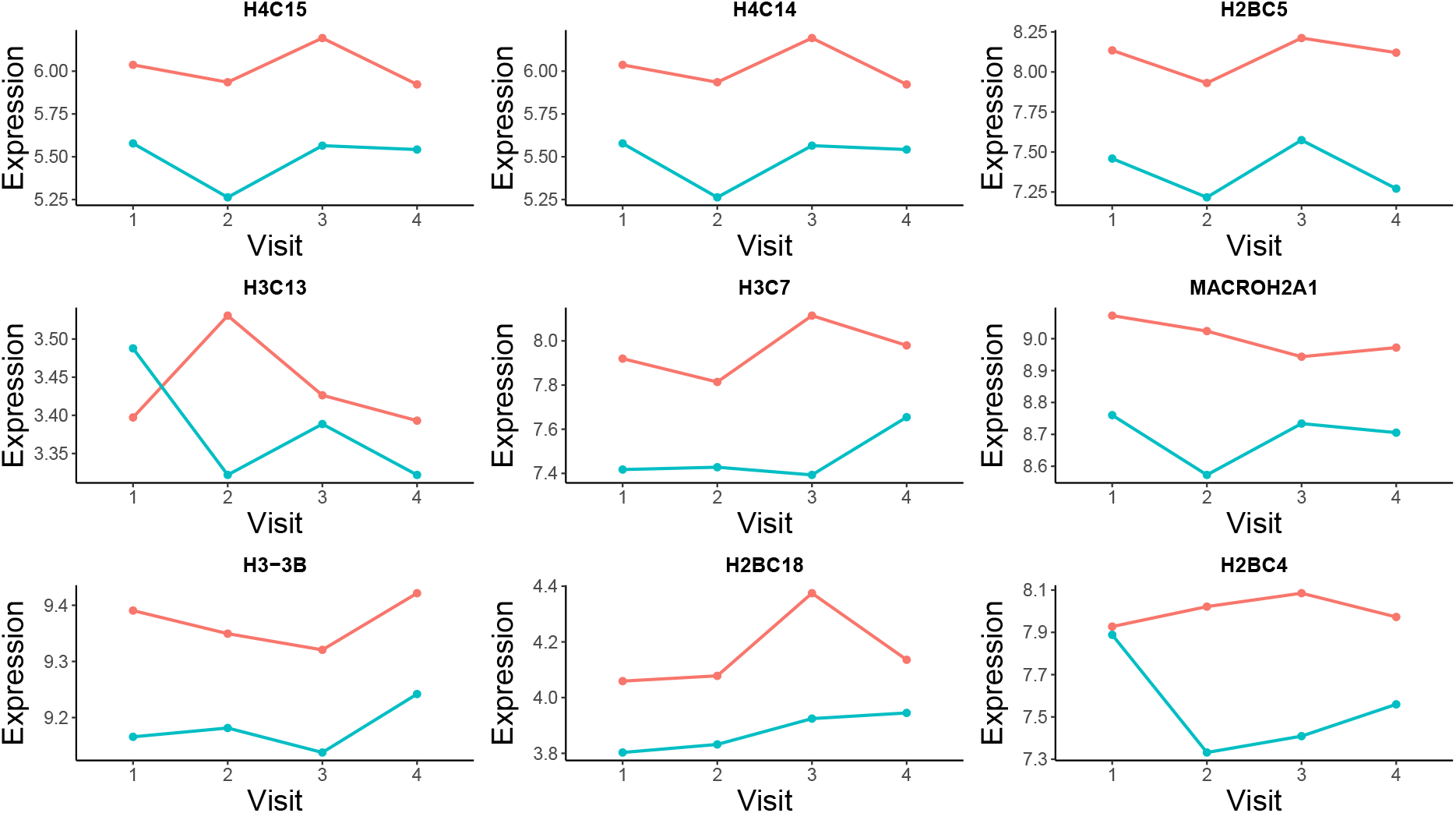
Differential expression of a module of nine histone genes. The mean expression profiles of each gene for 4 clinical visits are compared across 2 groups: AA (shown in red) and H (blue). Each gene’s symbol is shown above its profiles.

**Table 1:**
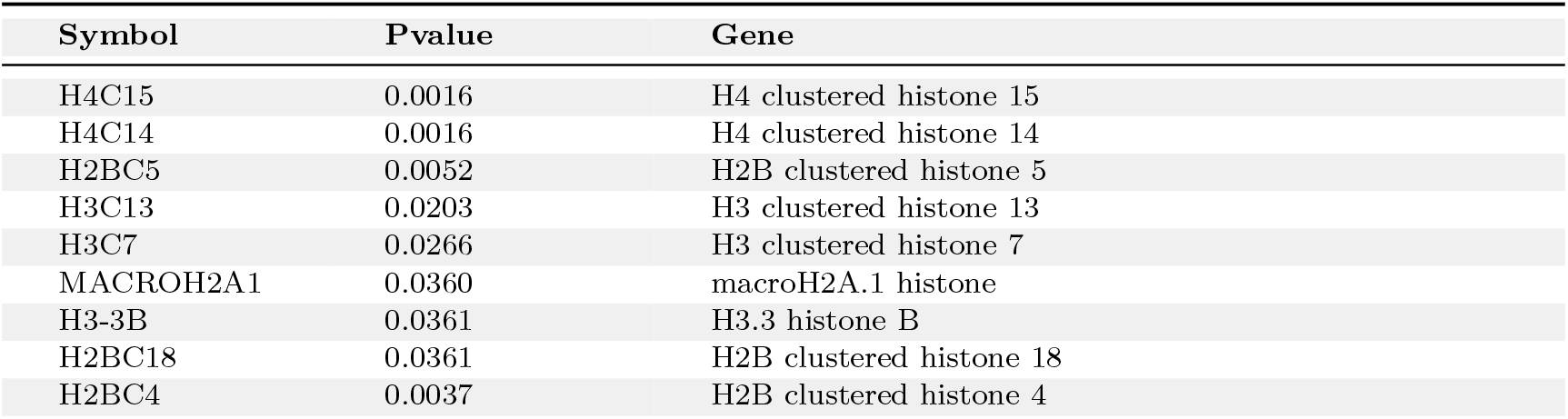
A differentially expressed module of histone genes associated with SLE.

Moreover, our analysis revealed profiles of gene expression in two other complementary scenarios. First, we identified genes that are mildly statistically significant and yet found in the literature to be involved in SLE. To illustrate, while the mean profiles of the C1QB gene are distinct in Figure 3a, it is weakly differentially expressed across the multiple groups and clinical visits, with a p-value of 0.0435. The gene encodes for the *C1qB* chain, a component of the complement system that functions as part of the immune system (Walport *et al*., 1998). Its deficiency is associated with a high probability of developing SLE as well as the severity and outcomes of the disease especially due to the impaired clearance of apoptotic debris as a result (Pickering *et al*., 2000). We listed the SLE-rlated phenotypes in HPO that are associated with C1QB in Table 2.

**Figure 3:**
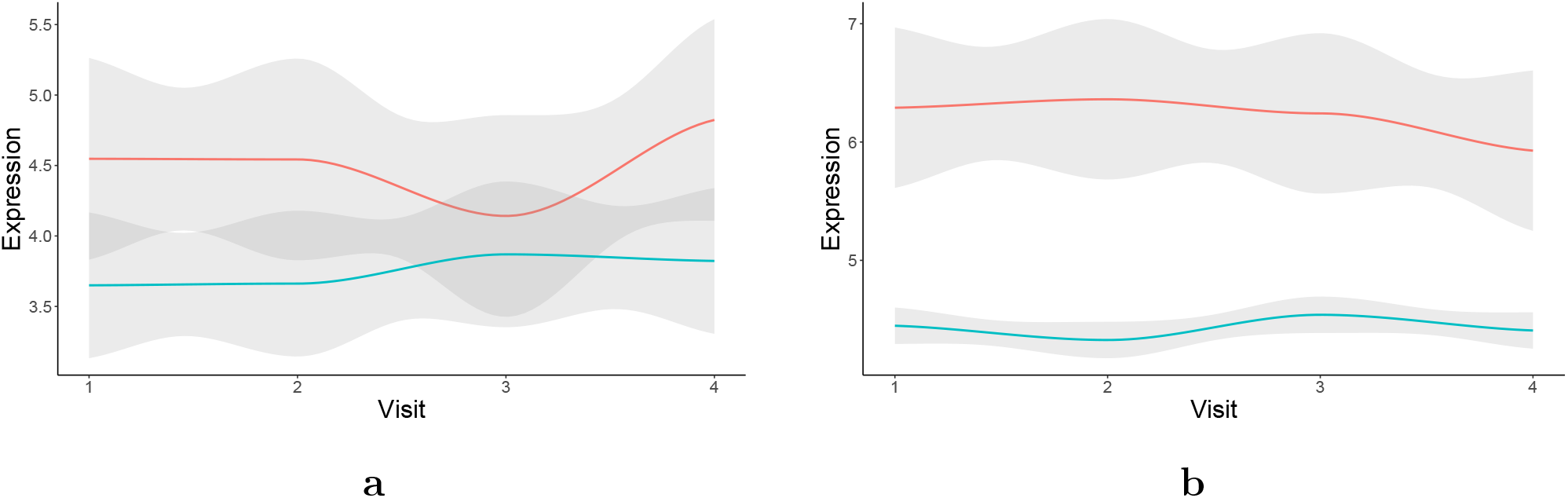
Differential expression of genes: (a) C1QB and (b) SOS1. The smoothed mean expression profiles and corresponding standard error (grey bands) of each gene for 4 clinical visits are compared across 2 groups: AA (red) and H (blue).

**Table 2:**
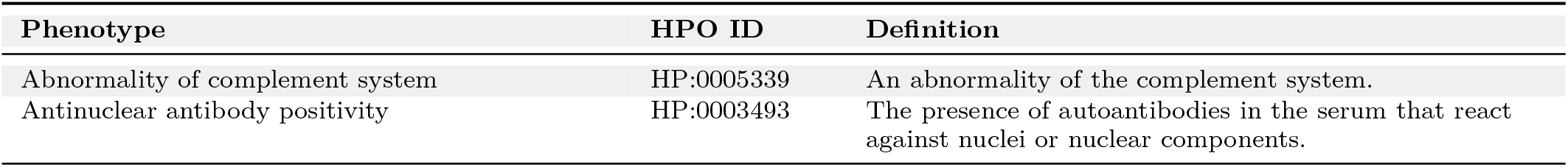
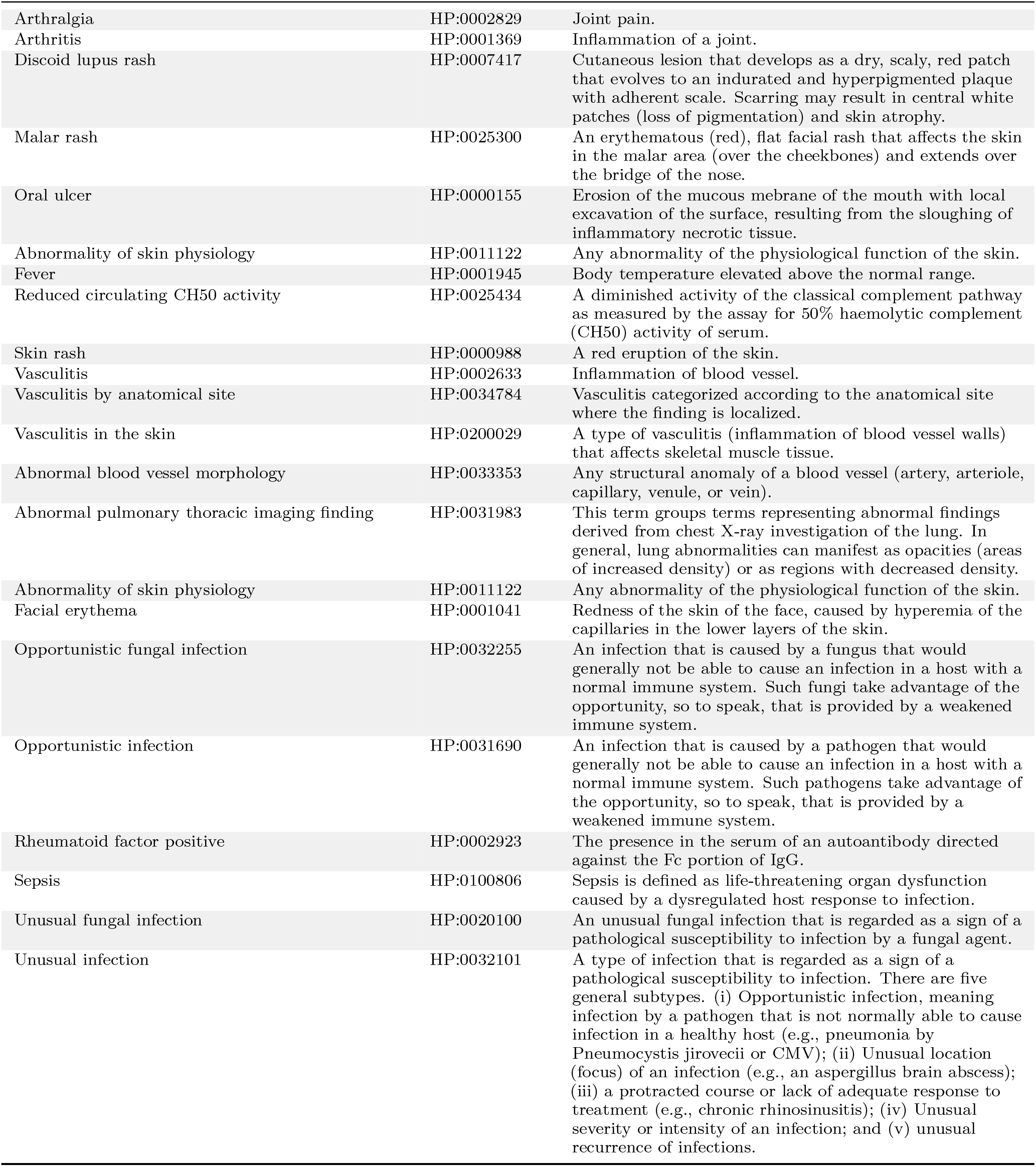
SLE-relevant phenotypes in HPO associated with C1QB gene expression.

Conversely, SOS1 is a gene that we found to be very strongly differentially expressed (p-value 0.0001) during the progression of SLE but it is not known to be directly linked to the disease. SOS1 encodes for a protein that acts as a master switch in the Ras/MAPK pathway, which shows a close association with autoimmune diseases (Zhang *et al*., 2025). By noting the differential expression of its profiles across the AA and H groups, as shown in Figure 3b, and the high count of SLE-relevant phenotypes **–** curated by HPO **–** that are linked to this gene and listed in Table 3, we think SOS1 is a highly suitable candidate biomarker for further investigation in this context.

**Table 3:** SLE-relevant phenotypes in HPO associated with SOS1 gene expression.

| Phenotype | HPO ID | Definition |
| --- | --- | --- |
| Prolonged partial thromboplastin time | HP:0003645 | Increased time to coagulation in the partial thromboplastin time (PTT) test, a measure of the intrinsic and common coagulation pathways. Phospholipid, and activator, and calcium are mixed into an anticoagulated plasma sample, and the time is measured until a thrombus forms. |
| Abnormal blood vessel morphology | HP:0033353 | Any structural anomaly of a blood vessel (artery, arteriole, capillary, venule, or vein). |
| Abnormal conjugate eye movement | HP:0000549 | Any deviation from the normal motor coordination of the eyes that allows for bilateral fixation on a single object. |
| Abnormal heart valve morphology | HP:0001654 | Any structural abnormality of a cardiac valve. |
| Abnormal heart valve physiology | HP:0031653 | Any functional abnormality of a cardiac valve. |
| Abnormal pulmonary valve morphology | HP:0001641 | Any structural abnormality of the pulmonary valve. |
| Abnormal pulmonary valve physiology | HP:0031654 | Any functional anomaly of the pulmonary valve. |
| Abnormality of fluid regulation | HP:0011032 | An abnormality of the regulation of body fluids. |
| Abnormality of joint mobility | HP:0011729 | An abnormality in the range and ease of motion of joints across their normal range. |
| Abnormality of skin pigmentation | HP:0001000 | An abnormality of the pigmentation of the skin. |
| Abnormality of the abdominal organs | HP:0002012 | An abnormality of the viscera of the abdomen. |
| Abnormality of the amniotic fluid | HP:0001560 | Abnormality of the amniotic fluid, which is the fluid contained in the amniotic sac surrounding the developing fetus. |
| Abnormality of the orbital region | HP:0000315 | No definition found. |
| Abnormality of the pulmonary vasculature | HP:0004930 | No definition found. |
| Abnormality of thrombocytes | HP:0001872 | An abnormality of platelets. |
| Abnormality of upper limb joint | HP:0009810 | No definition found. |
| Hypertrophic cardiomyopathy | HP:0001639 | Hypertrophic cardiomyopathy (HCM) is defined by the presence of increased ventricular wall thickness or mass in the absence of loading conditions (hypertension, valve disease) sufficient to cause the observed abnormality. |

## 4. Discussion

In this study, we demonstrated how a combined approach driven by data (HPO) and model (GCM) could yield differential gene expression profiles useful for inferring clinical phenotypes that might arise during the course of a heterogeneous disease. This is especially relevant to progression of SLE in which various phenotypes can dynamically and differentially occur across different racial groups. SLE is commonly referred to as a “cruel mystery” due to its wide range of symptoms that unpredictably varies from one patient to another, difficulty in diagnosis, and the potential to cause severe complications affecting the heart, lungs, and kidneys. SLE afflicts estimated 1.5 million people in the U.S. alone and 5 million worldwide. Typically, cases of pediatric SLE have more severe disease and damages in different organs (Lo, 2018). Hence, it is important to develop a statistical model that can infer dynamically emerging phenotypes based on longitudinal transcriptomic signals specific to populations.

Our study demonstrates all of these advantages. The GCM framework brought together (a) longitudinal immuno-monitoring data across races, which yielded (b) significantly differentially expressed gene signatures during disease progression, and these were useful for (c) systematic molecular characterization of the SLE phenotypes based on HPO annotations. Thus, it provided us with crucial insights into a variety of clinical phenotypes of pediatric lupus via detection of underlying transcriptomic profiles that were found to differ during disease progression between AA and H, the two population groups significantly affected by SLE in the U.S.

Abnormal histone modifications are known to play a major role in disease severity and outcomes of SLE (Araki and Mimura, 2017), e.g., via dysregulation of lupus CD4+ T cells (Hu *et al*., 2008). In this study, GCM identified a module of histone genes that are associated the production of NETs, which are released by dying neutrophils and whose role in SLE has been a topic of intense research in the past decade (Wigerblad and Kaplan, 2022). Here, we demonstrated their distinct profiles in the AA and H groups that determine SLE outcomes during race-specific progressions of pediatric SLE, which is less commonly studied in the literature.

We understand that our study has certain limitations such as the small sample size. In addition, no comparative analysis with other methods is provided. We plan to use larger studies for detailed simulations to compare with more complex linear mixed effects models in future work. Due to the lack of further datasets, cross-validation or replication studies, our findings are exploratory rather than confirmatory. Yet, notable molecular signatures identified as significantly differentially expressed showed clear patterns of consistency over the course of disease progression and is, thus, a demonstration of the effectiveness of our GCM approach to address this important problem.

